# Coincident glutamatergic depolarizations enhance GABA_A_ receptor-dependent Cl^-^ influx in mature and suppress Cl^-^ efflux in immature neurons

**DOI:** 10.1101/2020.08.18.255323

**Authors:** Aniello Lombardi, Peter Jedlicka, Heiko J. Luhmann, Werner Kilb

## Abstract

The impact of GABAergic transmission on neuronal excitability depends on the Cl^−^-gradient across membranes. However, the Cl^−^-fluxes through GABA_A_ receptors alter the intracellular Cl^−^ concentration ([Cl^−^]_i_) and in turn attenuate GABAergic responses, a process termed ionic plasticity. Recently it has been shown that coincident glutamatergic inputs significantly affect ionic plasticity. Yet how the [Cl^−^]_i_ changes depend on the properties of glutamatergic inputs and their spatiotemporal relation to GABAergic stimuli is unknown. To investigate this issue, we used compartmental biophysical models of Cl^−^ dynamics simulating either a simple ball-and-stick topology or a reconstructed immature CA3 neuron. These computational experiments demonstrated that glutamatergic co-stimulation enhances GABA receptor-mediated Cl^−^ influx at low and attenuates or reverses the Cl^−^ efflux at high initial [Cl^−^]_i_. The size of glutamatergic influence on GABAergic Cl^−^-fluxes depends on the conductance, decay kinetics, and localization of glutamatergic inputs. Surprisingly, the glutamatergic shift in GABAergic Cl^−^-fluxes is invariant to latencies between GABAergic and glutamatergic inputs over a substantial interval. In agreement with experimental data, simulations in a reconstructed CA3 pyramidal neuron with physiological patterns of correlated activity revealed that coincident glutamatergic synaptic inputs contribute significantly to the activity-dependent [Cl^−^]_i_ changes. Whereas the influence of spatial correlation between distributed glutamatergic and GABAergic inputs was negligible, their temporal correlation played a significant role. In summary, our results demonstrate that glutamatergic co-stimulation had a substantial impact on ionic plasticity of GABAergic responses, enhancing the destabilization of GABAergic inhibition in the mature nervous systems, but suppressing GABAergic [Cl^−^]_i_ changes in the immature brain. Therefore, glutamatergic shift in GABAergic Cl^−^-fluxes should be considered as a relevant factor of short term plasticity.

**Author Summary:** Information processing in the brain requires that excitation and inhibition are balanced. The main inhibitory neurotransmitter in the brain is gamma-amino-butyric acid (GABA). GABA actions depend on the Cl^−^-gradient, but activation of ionotropic GABA receptors causes Cl^−^-fluxes and thus reduces GABAergic inhibition. Here, we investigated how a coincident membrane depolarization by excitatory, glutamatergic synapses influences GABA-induced Cl^−^-fluxes using a biophysical compartmental model of Cl^−^ dynamics, simulating either simple or realistic neuron topologies. We demonstrate that glutamatergic co-stimulation directly affects GABA-induced Cl^−^-fluxes, with the size of glutamatergic effects depending on the conductance, the decay kinetics, and localization of glutamatergic inputs. We also show that the glutamatergic shift in GABAergic Cl^−^-fluxes is surprisingly stable over a substantial range of latencies between glutamatergic and GABAergic inputs. We conclude from these results that glutamatergic co-stimulation alters GABAergic Cl^−^-fluxes and in turn affects the strength of GABAergic inhibition. These coincidence-dependent ionic changes should be considered as a relevant factor of short term plasticity in the CNS.

## 1. Introduction

Information processing between single neurons is carried out via synaptic contacts, that use different neurotransmitters acting on pre- and postsynaptic receptors. The two most important neurotransmitters in the mammalian brain are glutamate and γ-amino butyric acid (GABA), which in general exert an excitatory and inhibitory action in the postsynaptic cell, respectively [1]. Beside regulating the excitability of neuronal circuits, GABA is also required for the control of sensory integration, regulation of motor functions, generation of oscillatory activity, and neuronal plasticity [2]. The responses to GABA are mediated by metabotropic GABA_B_ receptors and byionotropic GABA_A_ receptors, which are ligand-gated anion-channels with a high Cl^−^ permeability and a partial HCO3^-^ permeability [1]. The GABA actions therefore depend mainly on the Cl^−^ gradient across the neuronal plasma membrane [1,3]. The typical inhibitory action of GABA requires a low intracellular Cl^−^ concentration ([Cl^−^]_i_), which is typically maintained by the action of a Cl^−^ extruder termed KCC2 [3]. During early development [Cl^−^]_i_ is maintained at high levels by active Cl^−^ accumulation via the Na^+^-dependent K^+^-2Cl^−^-Symporter NKCC1, rendering GABA responses depolarizing and under certain conditions excitatory [4–6]. Such excitatory GABAergic responses contribute to the generation of spontaneous neuronal activity, which is essential for the correct maturation of nervous systems [7–9].

However, the Cl^−^-fluxes through GABA_A_ receptors, which underlie GABAergic currents, change [Cl^−^]_i_ and thus temporarily affect the amplitude of subsequent GABAergic responses, a process termed ionic-plasticity [3,10,11]. Such activity-dependent [Cl^−^]_i_ transients have been observed in various neurons [12–19]. Ionic plasticity plays an important role for physiological functions [20–22] as well as pathophysiological processes [23,24]. Theoretical assumptions and computational studies indicate that the amount and duration of [Cl^−^]_i_ ionic plasticity directly depends on the relation between Cl^−^ influx and the capacity of Cl^−^ extrusion systems [3,24–28]. In addition, the size and geometrical structure of the postsynaptic compartments critically affect the magnitude, duration and dimensions of [Cl^−^]_i_ changes upon GABAergic activation [27,29]. Further analyses also revealed that the membrane resistance, the kinetics of GABAergic responses and the stability of bicarbonate gradients affects the magnitude and duration of [Cl^−^]_i_ changes [12,27,30].

Another factor that directly influences the amount of GABA_A_ receptor-mediated Cl^−^ fluxes is a coincident membrane depolarization [11,27,31]. Accordingly, recent in-vitro and in-silico studies demonstrated that coincident glutamatergic depolarization profoundly augments the GABA_A_ receptor mediated Cl^−^ fluxes [32,33]. However, to our knowledge it has not been analyzed how the interdependency between glutamatergic and GABAergic inputs affects the magnitude and the spatiotemporal properties of activity dependent [Cl^−^]_i_ changes.

In order to determine the influence of coincident glutamate stimulation on GABA_A_ receptor induced [Cl^−^]_i_ transients, we utilized a computational model of [Cl^−^]_i_ homeostasis in the NEURON environment [25,30]. Using a simple ball and stick geometry, we were able to show that the conductance and decay kinetics of glutamatergic responses directly affects the size of GABAergic [Cl^−^]_i_ changes, with a complex spatiotemporal dependency between glutamatergic and GABAergic inputs. Furthermore, we employed the model to uncover the contribution of coincident glutamatergic activity on the activity-dependent [Cl^−^]_i_ transients during simulated giant-depolarizing potentials (GDPs) [17,30]. GDPs represent correlated spontaneous network events crucial for the development of neuronal circuits [34–36]. These simulations demonstrate for the first time that coincident glutamatergic stimulation with realistic parameters substantially modifies the GABA_A_ receptor-induced [Cl^−^]_i_ changes.

## 2. Results

In order to analyze how co-activation of depolarizing neurotransmitter receptors affects the [Cl^−^]_i_ transients evoked by GABA_A_ receptor activation, we used a previously established biophysical model of Cl^−^ dynamics in the NEURON environment [17,25]. The model allows for realistic simulations of activity-dependent transmembrane Cl^−^ fluxes, intracellular Cl^−^ diffusion and corresponding intracellular Cl^−^ accumulation or depletion (see Methods). In a first step we computed the GABA-induced [Cl^−^]_i_ changes in a ball and stick model with a single dendrite (Fig. 1a), in order to enable a detailed mechanistic understanding of the interaction between a single or a small group of GABA_A_ receptor-mediated and glutamate receptor-mediated synaptic inputs (Fig. 1 - 6). Subsequently we utilized a model of a reconstructed CA3 pyramidal neuron [17,30] stimulated by a large number of inputs to compute a more realistic estimation of how the GABA_A_ receptor-mediated [Cl^−^]_i_ transients in neurons are influenced by correlated co-activation of glutamate receptors (Fig. 7 - 9).

**Figure 1.**
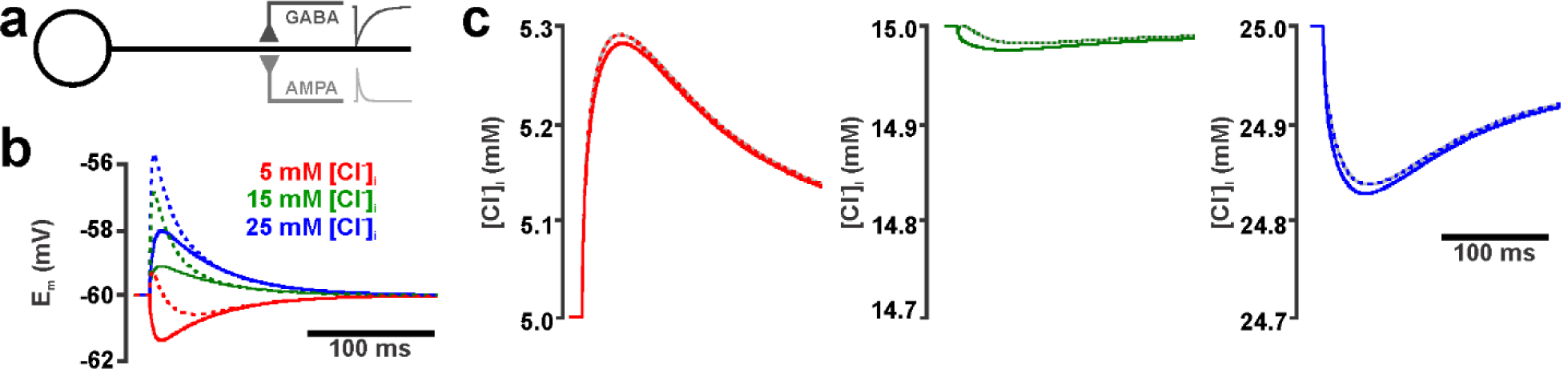
Effect of AMPA co-activation on GABA_A_ receptor mediated [Cl^−^]_i_ transients for a single GABAergic and glutamatergic input in the ball-and-stick neuronal model. (a) Schematic illustration of the compartmental model. Both GABA and AMPA synapse were located in the middle of the dendrite. (b) Membrane potential (E_m_) changes induced by a single GABAergic stimulation (g_GABA_ = 0.789 nS, τ_GABA_ = 37 ms) at 5 mM (red), 15 mM (green) and 25 mM [Cl^−^]_i_ (blue). Solid lines represent the effect upon an isolated GABAergic stimulation, while dashed lines represent responses upon AMPA co-stimulation (g_AMPA_ = 0.305 nS, τ_AMPA_ = 11 ms). (c) [Cl^−^]_i_ transients induced by the isolated GABAergic stimulation (solid lines) or the GABA/AMPA co-stimulation (dashed lines). Parameters as in (b). Note that under this conditions AMPA co-activation slightly enhances the [Cl^−^]_i_ increase at 5 mM, while it slightly attenuates the [Cl^−^]_i_ decline at higher [Cl^−^]_i_ concentrations.

### 2.1 Effect of GABA and AMPA co-activation for a weak focal synaptic activation

First we analyzed the effect of glutamate receptor co-activation on the GABA_A_ receptor-induced [Cl^−^]_i_ transients. It is important to emphasize that simulations of single or a low number of synaptic inputs necessarily leads to small absolute effects of glutamatergic activation on GABAergic [Cl^−^]_i_ or membrane voltage (E_m_) changes (Fig. 1 - 6), but nevertheless allowed for systematic parameter scans important for the detailed understanding of glutamatergic effects (see below). For quantitatively stronger effects during more realistic GABAergic and glutamatergic distributed input activation see Sections 2.4 and 2.5.

Simulation of a single GABA synapse using parameters determined in-vitro for spontaneous GABAergic postsynaptic currents (PSCs) (g_GABA_ = 0.789 nS, τ_GABA_ = 37 ms) [17], induced [Cl^−^]_i_ changes that depended on the initial [Cl^−^]_i_ (Fig. 1b, c), as shown before [17]. At low initial [Cl^−^]_i_ the inwardly directed driving force (DF_GABA_) caused a Cl^−^-influx and thus a membrane hyperpolarization. At higher initial [Cl^−^]_i_ of 25 mM the outwardly directed DF_GABA_ caused a Cl^−^-efflux and thus a membrane depolarization, while at an initial [Cl^−^]_i_ of 15 mM, which is close to the passive [Cl^−^]_i_ at -60 mV, only a slight [Cl^−^]_i_ efflux was induced (Fig. 1b, c). The simultaneous co-activation with a simulated single glutamate synapse, using parameters that were determined in-vitro for spontaneous glutamatergic PSCs [17]) and resemble the properties of the AMPA subtype of glutamate receptors *(g*_*AMPA*_ *= 0*.*305 nS, τ*_*AMPA*_ *= 11 ms)*, imposed an additional depolarizing drive to the E_m_ responses (Fig. 1b, dashed lines). Thereby it caused slight changes in the activity-dependent [Cl^−^]_i_ transients towards higher concentration, thus enhancing the [Cl^−^]_i_ increase at low initial [Cl^−^]_i_ and attenuating the [Cl^−^]_i_ decrease at higher initial [Cl^−^]_i_ (Fig. 1c, dashed lines).

Since physiological and pathophysiological activity is typically characterized by neurotransmitter release from several release sites [37,38], we next analyzed the effect of varying g_AMPA_ on the GABA_A_ receptor -mediated [Cl^−^]_i_ transient. These simulations revealed that without co-stimulation GABA_A_ receptor induced Cl^−^ fluxes depended only on the initial [Cl^−^]_i_ (Fig. 2a). In contrast, the Cl^−^ fluxes were systematically shifted towards Cl^−^ influx in the presence of AMPA receptor mediated inputs (Fig. 2a). At a g_AMPA_ of 305 nS and 30.5 nS (corresponding to 100x and 1000x glutamatergic PSCs) a consistent Cl^−^ influx was induced even at high [Cl^−^]_i_. At a g_AMPA_ of 3.05 nS (Fig. 2a, blue line) and 0.305 nS (Fig. 2a, pink line) bimodal effects, consisting of an initial influx followed by an efflux, were observed within a limited [Cl^−^]_i_ range (Fig. 2b, c). These biphasic responses were caused by the fact that the membrane depolarization only temporarily exceeds E_Cl_ (Fig. 2b, c). The biphasic effects are displayed by the two lines representing in- and efflux for 3.05 nS (blue line) and 0.305 nS (pink line) in Fig. 2a.

**Figure 2.**
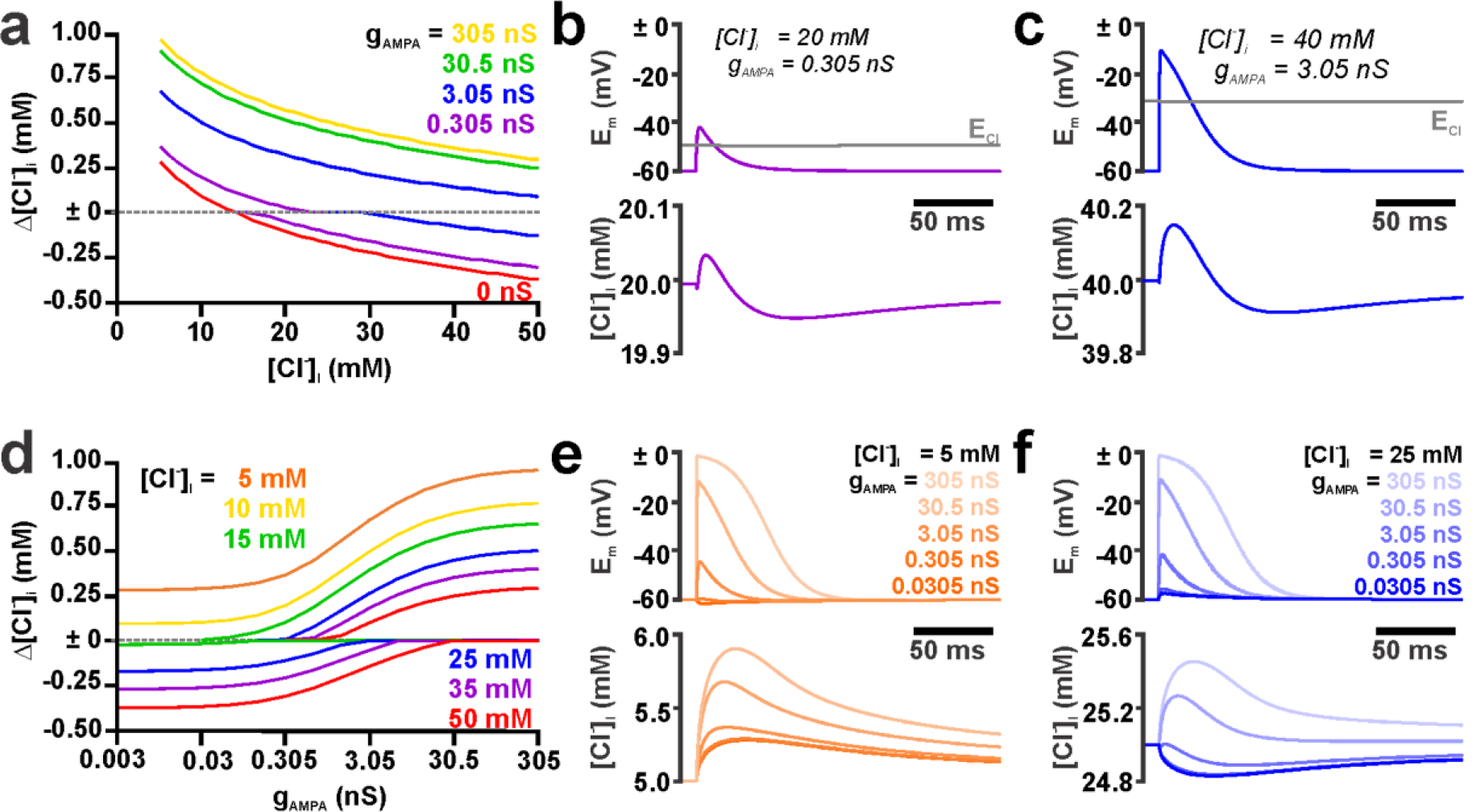
Effect of g_AMPA_ on the [Cl^−^]_i_ transients induced by GABA_A_ and AMPA receptor co-activation. (a) GABA_A_ receptor-induced [Cl^−^]_i_ changes simulated at different initial [Cl^−^]_i_ in the absence (red trace) and the presence of AMPA receptor co-activation with different g_AMPA_ as indicated in the plot. The GABA synapse was stimulated with a g_GABA_ of 0.789 nS and a τ_GABA_ of 37 ms. For g_AMPA_ of 0.305 pS (purple lines) and 3.05 pS (blue lines) biphasic [Cl^−^]_i_ transients were observed (see b and c) and thus minimal and maximal [Cl^−^]_i_ were depicted in the graph for the same initial [Cl^−^]_i_ value. (b) Membrane depolarization (upper panel) and [Cl^−^]_i_ (lower panel) at a [Cl^−^]_i_ of 20 mM and a g_AMPA_ of 0.305 nS. Note the biphasic [Cl^−^]_i_ transient and that the Cl^−^ influx was limited to the interval when Em is above E_Cl_ (gray trace). (c) As in b but for 40 mM and g_AMPA_ of 3.05 nS. (d) g_AMPA_ dependency of the GABA_A_ receptors induced [Cl^−^]_i_ changes at different initial [Cl^−^]_i_ as indicated in the plot. Note the shift towards Cl-influx with increasing g_AMPA_. Similarly to a, when biphasic [Cl^−^]_i_ transients were observed, both minimal and maximal [Cl^−^]_i_ were depicted in the graph for the same g_AMPA_ value. (e) Membrane depolarization (upper panel) and [Cl^−^]_i_ (lower panel) at a [Cl^−^]_i_ of 5 mM and various g_AMPA_, as indicated in the plot. Note that increasing g_AMPA_ systematically enhanced Cl^−^ influx at this initial [Cl^−^]_i_. (f) (e) Membrane depolarization and [Cl^−^]_i_ at a [Cl^−^]_i_ of 25 mM and various g_AMPA_. Note that at this initial [Cl^−^]_i_ the Cl^−^ efflux was shifted towards Cl^−^ influx with increasing g_AMPA_. In all simulations g_GABA_ = 0.789 nS, τ_GABA_ = 37 ms and τ_AMPA_ = 11 ms.

A systematic analysis of the direction of Cl^−^ fluxes at various g_AMPA_ revealed that the GABA_A_ receptor mediated Cl^−^ fluxes show a logarithmic dependency on g_AMPA_, resulting in sigmoidal curves in a monologarithmic plot (Fig. 2d). Only at low [Cl^−^]_i_ concentrations, which permit a Cl^−^ influx even without AMPA co-stimulation, a monotonous increase in the Cl-fluxes were induced by increasing g_AMPA_ (Fig. 2d, e). When [Cl^−^]_i_ was above 13 mM (corresponding to E_Cl_ above -60 mV) addition of g_AMPA_ caused biphasic Cl-fluxes (Fig. 2d, f).

In summary, these results indicate that co-activation of AMPA receptors enhances Cl^−^ influx at low [Cl^−^]_i_ and attenuates Cl^−^ efflux at high [Cl^−^]_i_. with the possibility to even revert the latter to Cl^−^ influx. Already relatively low, physiological relevant g_AMPA_ values, representing 10-100 spontaneous postsynaptic events, are sufficient to substantially modulate GABA_A_ receptor mediated Cl^−^ fluxes.

### 2.2 Influence of τ_AMPA_ on GABA_A_ receptor induced [Cl^−^]_i_ transients

Next we analyzed the influence of the kinetics of coincident AMPA receptor activation on the GABA_A_ receptor-induced [Cl^−^]_i_ transients. For this purpose, we modified the decay time constant of the AMPA synapse (τ_AMPA_) using values between 1 and 100 ms. These simulations were performed under consideration of the experimentally determined values for GABA_A_ receptor-mediated inputs either at gAMPA of 0.305 pS (i.e. the experimentally determined value [17]) or at gAMPA of 3.05 nS (to estimate the effects of a small number of coincident colocalized glutamatergic inputs). As the depolarizing effect of the AMPA synapse strictly depends on τ_AMPA_ (Fig. 3a), for τ_AMPA_ values < 37 ms biphasic voltage responses were induced at low [Cl^−^]_i_ (Fig. 3a). In contrast, at high [Cl^−^]_i_ (where GABAergic responses are depolarizing) the peak depolarization was enhanced. For τ_AMPA_ values > 37 ms the depolarization was virtually saturated (Fig. 3b). In accordance with this observation the AMPA-mediated shift in the GABAergic [Cl^−^]_i_ transients was sensitive to τ_AMPA_ at values < 37 ms, while at values > 37ms only minimal additional effects were observed (Fig. 3c, d). These findings indicate that the kinetics of glutamatergic synaptic inputs have a consistent effect on activity dependent [Cl^−^]_i_ changes and that shorter glutamatergic postsynaptic potentials (PSPs) will attenuate the effect of AMPA on activity-dependent [Cl^−^]_i_ transients.

**Figure 3.**
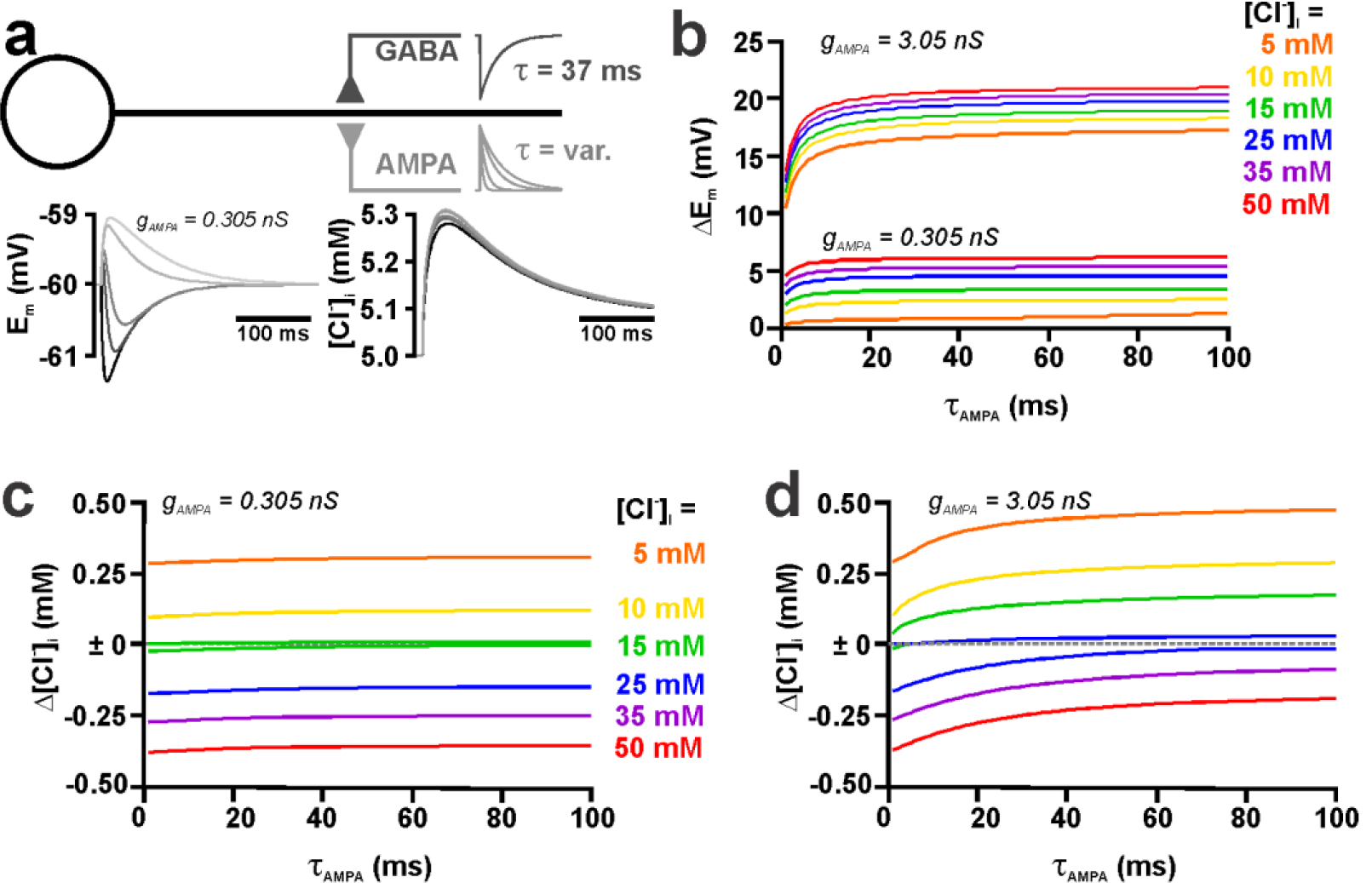
Effect of τ_AMPA_ on the [Cl^−^]_i_ transients induced by GABA_A_ and AMPA receptor co-stimulation. (a) Schematic illustration of the compartmental model. The GABA synapse (g_GABA_= 0.789 nS) was stimulated with a constant τ_GABA_ of 37 ms, while τ_AMPA_ was systematically varied. The traces below the scheme displays the voltage responses (left panel) and [Cl^−^]_i_ changes induced without (black traces) or with AMPA co-stimulation (g_AMPA_ = 0.305 nS) at τ_AMPA_ values of 5, 11, 37, and 50 ms (in ascending gray shades). Note the biphasic voltage responses at short τ_AMPA._. (b) Membrane potential changes induced by a AMPA/GABA co-stimulation at different τ_AMPA_ simulated for different [Cl^−^]_i_ (color coded). The upper traces were simulated using a g_AMPA_ of 0.305 nS, while the lower traces represent simulation with g_AMPA_ of3.05 nS. Note that the depolarization increased with larger τ_AMPA_. (c) [Cl^−^]_i_ transients induced by AMPA/GABA co-stimulation at different τ_AMPA_ simulated with different [Cl^−^]_i_ (color coded) for a g_AMPA_ of 0.305 nS. (d) As in c) but for a g_AMPA_ of 3.05 nA. Note that the [Cl^−^]_i_ changes were shifted towards diminished Cl^−^ efflux or enhanced Cl^−^ influx with larger τ_AMPA_.

### 2.3 Spatial and temporal constrains of AMPA receptor-mediated shift in GABAergic [Cl^−^]_i_ transients for a weak focal synaptic activation

To analyze how the spatial distance between the AMPA and the GABA synapse influences the GABA_A_ receptor-induced [Cl^−^]_i_ transients, we systematically moved the AMPA synapse along the dendrite from the somatic (0%) to the distal end (100%) (Fig. 4a, b) using the experimentally determined parameters for AMPA and GABA inputs (g_GABA_ = 0.789 nS, τ_GABA_ = 37 ms, g_AMPA_ = 0.305 nS, τ_AMPA_ = 11 ms [17]). These experiments revealed that the membrane potential change depends on the location of the AMPA synapse. The maximal amplitude of the positive shift in E_m_ induced by the AMPA co-stimulation decreased substantially at distant positions (Fig. 4b), due to the leak conductance shunting the synaptic currents (i.e. dendritic filtering). In accordance with these smaller depolarizing E_m_ shifts, the [Cl^−^]_i_ changes were slightly decreased with increasing distance between AMPA and GABA synapses (Fig. 4b). To quantify this effect, we determined the amount of additional [Cl^−^]_i_ changes induced by AMPA co-stimulation (Δ_G_[Cl^−^]_i_ = [Cl^−^]_i_ at AMPA/GABA co-stimulation -[Cl^−^]_i_ at GABA stimulation), while systematically enhancing the distance between both synapses. These simulations revealed an exponential decay of the AMPA effect of [Cl^−^]_i_ with a decay length constant (λ) of 268 µm (Fig. 4c). This is exactly the λ value determined for the attenuation of the peak voltage responses (Fig. 4d). These observations indicate that the distance between GABAergic and glutamatergic synapses is a relevant factor determining the effects of co-activation on the activity-dependent [Cl^−^]_i_ transients.

**Figure 4.**
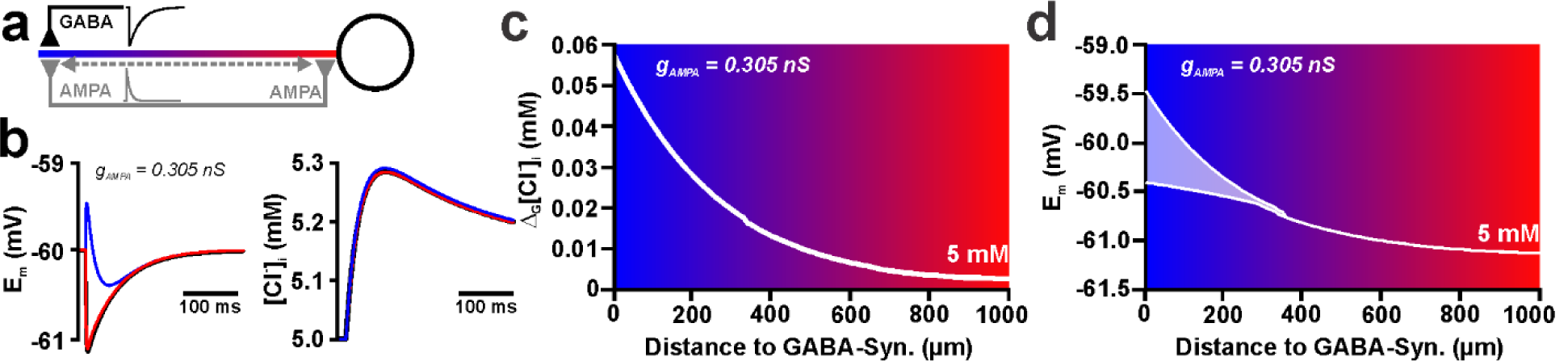
Effect of the spatial relation between individual GABA and AMPA synapses on the activity-dependent [Cl^−^]_i_ transients. (a) Schematic illustration of experimental conditions. Initial [Cl^−^]_i_ was 5 mM in these experiments. (b) Voltage responses (left panel) and [Cl^−^]_i_ changes (right panel) induced without (black traces) or with AMPA co-stimulation (g_AMPA_ = 0.305 nS) with the AMPA synapse located at the site of the GABA synapse (blue) or close to the soma (red). Note that the depolarization and [Cl^−^]_i_ responses are slightly diminished if the AMPA synapse was distant to the GABA synapse. (c) Effect of the distance between AMPA and GABA synapses on the additional [Cl^−^]_i_ influx induced by AMPA co-stimulation, as compared to a GABA stimulation without AMPA (Δ_G_[Cl^−^]_i_). Note the exponential decay of Δ_G_[Cl^−^]_i_ with increasing distance between GABA and AMPA synapses. (d) Effect of the distance between AMPA and GABA synapses on the resulting membrane depolarization. The upper and lower lines represent the maximal de- and hyperpolarizing effects for biphasic responses; above 400 µm only a monophasic hyperpolarization occurred.

To analyze how the temporal relation between the AMPA and the GABA mediated activity influences the GABA_A_ receptor-induced [Cl^−^]_i_ transients we next systematically varied the delay between the AMPA and the GABAergic stimulus from -49 ms (i.e. AMPA before GABA) to +100 ms (i.e. AMPA after GABA) and determined the impact of this latency shift on the [Cl^−^]_i_ transients (Fig. 5a, b). The parameters for AMPA and GABA inputs were identical to the parameters used before (g_GABA_ = 0.789 nS, τ_GABA_ = 37 ms, g_AMPA_ = 0.305 nS, τ_AMPA_ = 11 ms) and both synapses were located at the same position. These simulations revealed that the additional effect of AMPA co-stimulation on the [Cl^−^]_i_ transients (Δ_G_[Cl^−^]_i_) became maximal when GABA and AMPA stimulus were provided simultaneously (Fig. 5c). Surprisingly, this AMPA effect on Δ_G_[Cl^−^]_i_ remained stable for a latency of ≈20 ms, before it rapidly declined (Fig. 5c). To provide a mechanistic explanation for this plateau phase we next plotted the rate of [Cl^−^]_i_ changes versus the absolute value of the [Cl^−^]_i_ change (Fig. 5d). This phase plane plot illustrates that the trajectories of all AMPA stimulations with a latency between 0 and 20 ms converged with the trajectory of the 0 ms latency stimulus (purple line), thus reaching identical minimal [Cl^−^]_i_ values (obtained at the intersection with the y-axis value ∂[Cl^−^]_i_/∂t = 0 mM/s). Latencies between 22 and 34 ms provided a gradual decline in the minimal [Cl^−^]_i_. For latencies > 34 ms the additional AMPA-mediated reduction of the Cl^−^ -efflux was initiated after a minimal [Cl^−^]_i_ was reached, therefore these stimulations did not provide a [Cl^−^]_i_ decrease exceeding the [Cl^−^]_i_ decrease mediated by a pure GABA stimulation (black line).

**Figure 5.**
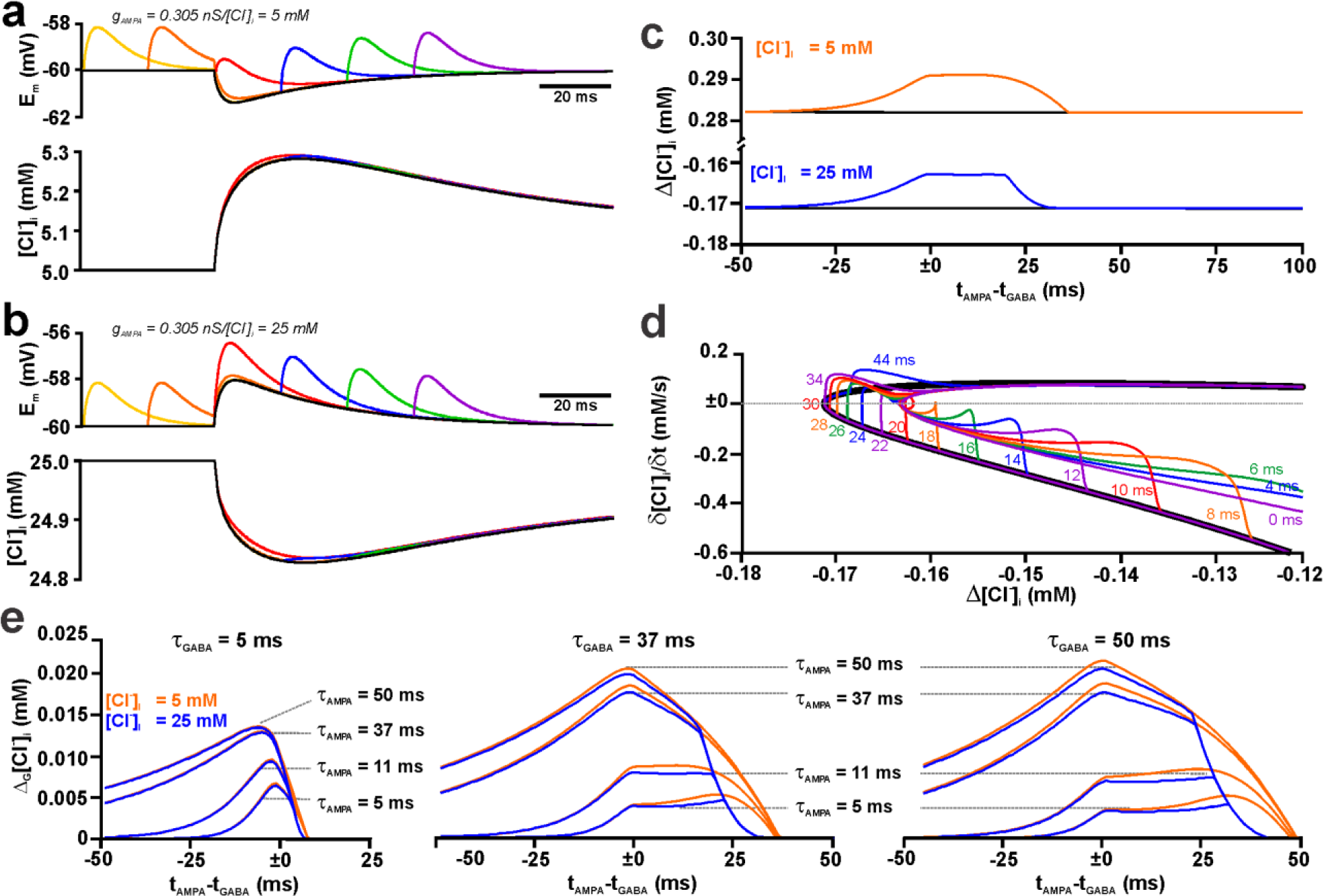
Effect of the temporal relation between single dendritic GABA and AMPA synapses on the activity-dependent [Cl^−^]_i_ transients. (a) Voltage (upper panel) and [Cl^−^]_i_ traces (lower panel) of [Cl^−^]_i_ changes induced by GABAergic stimulation without AMPA (black trace) or with AMPA stimulation at different latencies (colored traces) at an initial [Cl^−^]_i_ of 5 mM. (b) Same as in (a) for an initial [Cl^−^]_i_ of 25 mM. (c) Plot of the peak [Cl^−^]_i_ change at various latencies between GABA and AMPA inputs for an initial [Cl^−^]_i_ of 5 mM (orange trace) and 25 mM (blue trace). The black lines represent the [Cl^−^]_i_ change induced by stimulation of GABA synapses only. Note the obvious “plateau”-like phases in the [Cl^−^]_i_ changes that were additionally induced by AMPA costimulation (Δ_G_[Cl^−^]_i_). (d) Phase plane plot of the activity-dependent [Cl^−^]_i_ transients (efflux) at an initial [Cl^−^]_i_ of 25 mM. The black line represents the trajectory of a pure GABA stimulation. Note that all trajectories of 0 to 20 ms latency curves converged and crossed the 0 y-axis value (∂[Cl^−^]_i_ /∂t = 0 mM/s; dashed line) at a less negative x-axis value (Δ[Cl^−^]_i_) than the crossings of the >22 ms latency curves. This indicates that 0 – 20 ms latencies similarly and robustly decrease the peak GABA-induced efflux (i.e. 0 y-axis crossing of the pure GABA black line). See main text for further description. (e) Dependency of Δ_G_[Cl^−^]_i_ on the latency between AMPA and GABA stimulation simulated for different τ_GABA_ and τ_AMPA_. Note the plateau-like phases occurring for τ_GABA_ ≥ 37 ms at short τ_AMPA_.

To investigate how the duration of GABA and AMPA receptor-mediated currents influence this complex time course of [Cl^−^]_i_ transients, we next systematically varied τ_GABA_ (5 ms, 37, ms 50 ms) as well as τ_AMPA_ (5 ms, 11 ms, 37, ms 50 ms) and determined Δ_G_[Cl^−^]_i_ (Fig. 5e). These simulations revealed that, in accordance with the previous results, Δ_G_[Cl^−^]_i_ increased at longer τ_AMPA_ for all three τ_GABA_ tested. While at a short τ_GABA_ of 5 ms no plateau phase was observed, such a plateau occurred for τ_GABA_ of 37 ms and 50 ms with short τ_AMPA_ values of 5 ms or 11 ms (Fig. 5e). However, in contrast to our hypothesis that such a plateau phase occurred when τ_AMPA_ was shorter than τ_GABA,_ for a τ_GABA_ ≥ 37 ms and τ_AMPA_ ≥ 37 ms Δ_G_[Cl^−^]_i_ steadily declined after the maximal value was obtained at simultaneous AMPA/GABA stimulation. Another finding of these simulations that appears counter-intuitive is the fact that for short τ_AMPA_ of 5 ms and 11 ms the peak Δ_G_[Cl^−^]_i_ decreases when τ_GABA_ increased from 5 ms to 37 ms (Fig. 5e).

To investigate these issues in detail, we next systematically varied τ_GABA_ between 5 and 50 ms using fixed τ_AMPA_ values of 5, 11, and 37 ms and determined Δ_G_[Cl^−^]_i_.at different AMPA/GABA latencies. In these simulations we were able to identify a complex dependency between Δ_G_[Cl^−^]_i_ and the relation between τ_GABA_ and τ_AMPA_ (Fig. 6a). For τ_AMPA_ of 5 and 11 ms the influence of the AMPA/GABA latency on Δ_G_[Cl^−^]_i_ changed from a “peak”-like pattern to a “plateau”-like phase, in which the maximal Δ_G_[Cl^−^]_i_ remained rather constant for a progressively longer latency interval between AMPA and GABA stimulation. For τ_AMPA_ of 37 ms this “plateau”-like phase was replaced by a phase with linearly decreasing [Cl^−^]_i_ shifts (Fig. 6a). In any way, the maximal AMPA-dependent [Cl^−^]_i_ shift was observed at τ_GABA_ of 11 ms, 18 ms and 47 ms for τ_AMPA_ of 5 ms, 11 ms and 37 ms, respectively (red plots in Fig. 6a), reproducing the observation that prolonging τ_GABA_ can reduce Δ_G_[Cl^−^]_i_. From these results it can be concluded that the effect of AMPA co-stimulation has a complex influence on Δ_G_[Cl^−^]_i_ and that Δ_G_[Cl^−^]_i_ is maximal when τ_GABA_ is slightly larger than τ_AMPA_.

**Figure 6.**
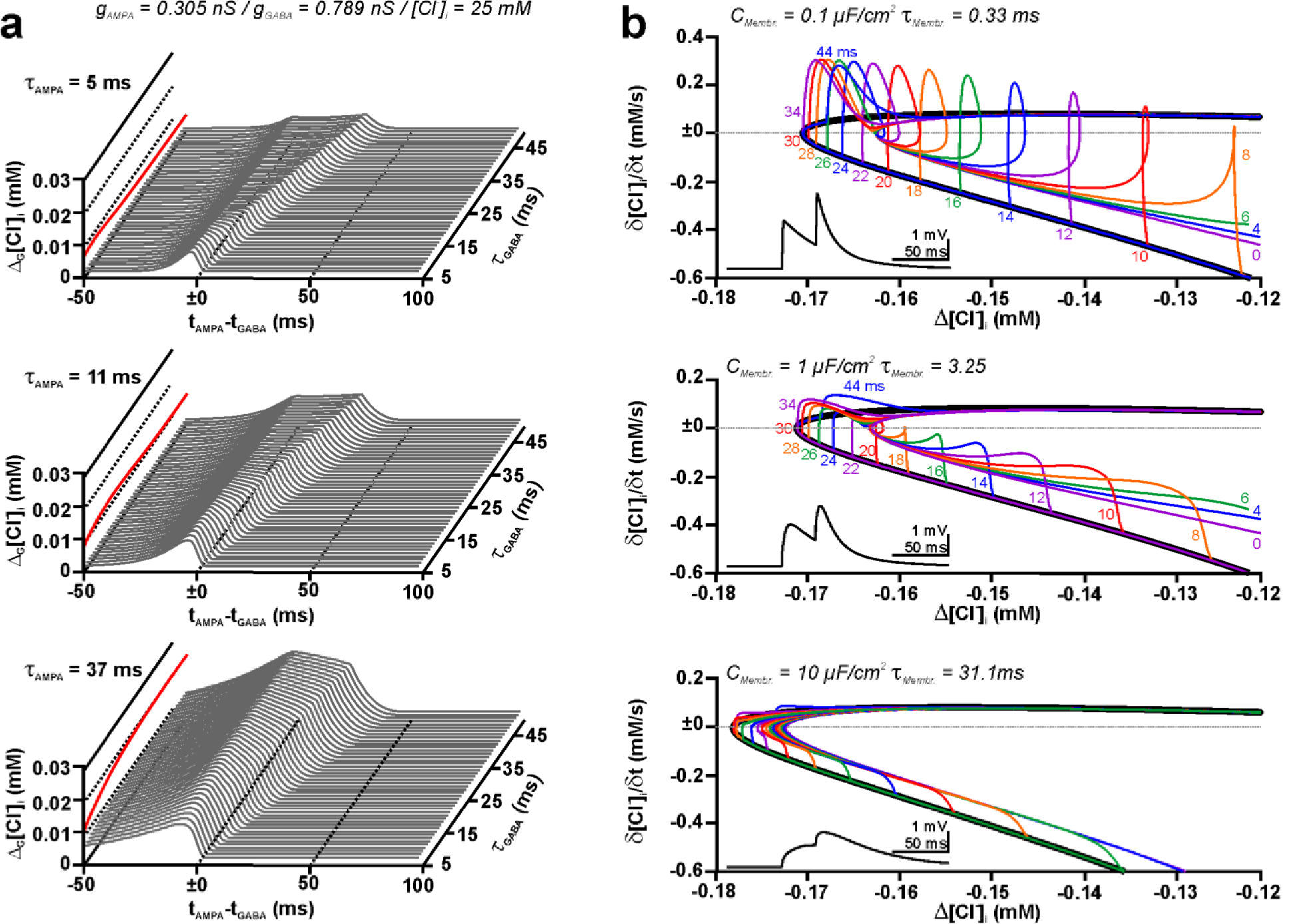
Effect of the relation between kinetics of individual dendritic GABA and AMPA synapses on the activity-dependent [Cl^−^]_i_ transients at different temporal relations between both synapses. (a) Amplitude of the additional [Cl^−^]_i_ influx induced by AMPA co-stimulation (compared to a GABA stimulation without AMPA) at different intervals between AMPA and GABA activation. Simulations were performed for 3 different τ_AMPA_ of 5 ms (top panel), 11 ms (middle panel), and 37 ms (lower panel) with τ_GABA_ varying between 5 ms and 50 ms. Note the evolution of a “plateau-like” phase with increasing τ_GABA_. The peak [Cl^−^]_i_ change was displayed as red trace on the left face of the plots. Note that for τ_AMPA_ of 5 ms and 11 ms, increasing τ_GABA_ above 10 ms and 17 ms reduced the [Cl^−^]_i_ shift induced by AMPA co-stimulation. (b) Effect of the membrane time constant on the trajectories of [Cl^−^]_i_ transients in a phase-plane plot. The black lines represent the [Cl^−^]_i_ change induced by stimulation of GABA synapses only. Typical voltage deflections are displayed in the insets. Note that decreasing the membrane time constant (upper panel) resulted in a comparable convergence of the trajectories towards the 0 ms latency (purple line) and GABA only conditions (black lines), despite the sharper trajectories. In contrast, after prolonging the membrane time constant (lower panel) the trajectories did not converge to the 0 ms latency condition at the intersection with ∂[Cl^−^]_i_ /∂t = 0 mM/s (dashed line).

Next, we investigated how the time course of the voltage deflections contribute to this complex dependency. For this purpose, we used identical stimulation parameters (g_GABA_ = 0.789 nS, τ_GABA_ = 37 ms, g_AMPA_ = 0.305 nS, τ_AMPA_ = 11 ms, initial [Cl^−^]_i_ = 25 mM) and modified the time course of the voltage traces by changing the membrane time constant (Fig. 6b). These simulations revealed that speeding up the kinetics of AMPA and GABA-receptor dependent voltage responses had only a minor effect on the observed [Cl^−^]_i_ changes (Fig. 6b, upper panel). The trajectories at different latencies converge virtually in the same manner as at physiological conditions (Fig. 6b, middle panel). On the other hand, a substantial prolongation of the membrane time constant slowed the kinetics of both AMPA and GABA-receptor dependent depolarizations (Fig. 6b, lower panel) and led to significantly different behavior or [Cl^−^]_i_ changes: The delayed GABAergic depolarization increased Δ[Cl^−^]_i_ when GABAergic input was stimulated alone (black trace). In addition, the maximum amount of Δ_G_[Cl^−^]_i_ (i.e. the difference between the 0 ms latency trajectory and the GABA only trajectory) was smaller. More importantly, the trajectories for latencies > 4 ms did no longer converge with the trajectory of simultaneous AMPA/GABA stimulation (purple line), which resulted in a steady decline of Δ_G_[Cl^−^]_i_ for all conditions in which AMPA was stimulated after the GABA input. In summary, these results indicate a complex interplay between the membrane time constant and the kinetics of AMPA- and GABA-mediated synaptic events. Only in cases in which the kinetics of the receptors dominate the time course of membrane responses, the complex, plateau-like dependency of Δ_G_[Cl^−^]_i_ on the latency between GABA and AMPA inputs were observed.

### 2.4 Influence of gAPMA and τAMPA on the GABAergic [Cl^−^]_i_ transients in a realistic neuronal model with multiple distributed synaptic inputs

To investigate in a more physiological setting how the interference between GABA_A_ and AMPA receptors influence the resulting [Cl^−^]_i_ transients, we implemented GABA and AMPA synapses in a morphologically and biophysically realistic model of an immature CA3 pyramidal neuron (Fig. 7a, b), using morphology and membrane parameters determined in *in vitro* experiments [17]. In the majority of simulations, we applied experimentally estimated correlated GABAergic and glutamatergic activity recorded during giant depolarizing potentials (GDPs, [17], see Fig. 7c). GDPs are highly relevant network events in the immature hippocampus that have been shown to cause substantial [Cl^−^]_i_ changes [8,17,35,39]. This computational model, which incorporates basic morphological and biophysical properties of hippocampal neurons, generated complex trajectories of [Cl^−^]_i_ within individual dendrites during a simulated GDP (Fig. 7d). These [Cl^−^]_i_ changes in the individual neurites are determined by Cl^−^-fluxes via GABA_A_ receptors, lateral Cl^−^ diffusion within the dendritic compartment and transmembrane Cl^−^-transport [25,29,33]. For a better quantification and display, the average [Cl^−^]_i_ over all nodes of all dendrites was calculated at each simulated interval (compare e.g. Fig. 7f).

**Figure 7.**
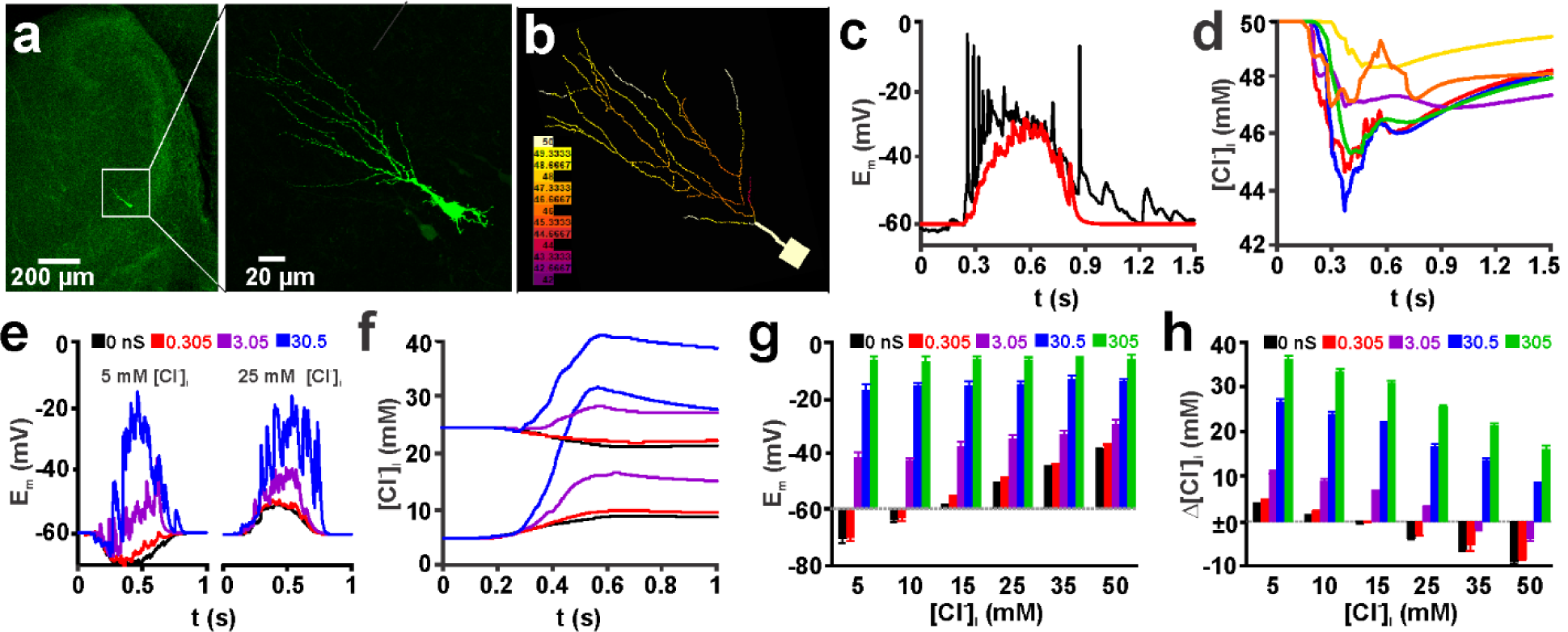
Effect of AMPA receptor co-stimulation on GABA_A_ receptor-induced [Cl^−^]_i_ transients in a realistic neuronal model stimulated with multiple spatially distributed synaptic inputs replicating GDP-like depolarizations (GDPs are experimentally observed giant depolarizing potentials in immature neurons). (a) Microfluorimetric image of a biocytin-filled CA3 pyramidal neuron stained during an electrophysiological recording. (b) Morphological representation of this neuron in the NEURON environment. The colors represent the actual [Cl^−^]_i_ during a simulated GDP. (c) Recorded (black-trace) and simulated (red trace) voltage deflections during a GDP with GABAergic synaptic inputs only. Note that no Hodgkin-Huxley spiking mechanisms were implemented in the model. (d) Time course of [Cl^−^]_i_ in the center node of 6 typical dendrites during this simulated GDP. (e) Typical voltage deflections during a GDP at an initial [Cl^−^]_i_ of 5 mM (left panel) and 25 mM (right panel) using different amounts of AMPA co-stimulation inputs as indicated by the color code. Note the substantial shift towards depolarized potentials at high g_AMPA_. (f) Time course of average dendritic [Cl^−^]_i_ at different initial [Cl^−^]_i_ and g_AMPA_ (as indicated in e). Note that at an initial [Cl^−^]_i_ of 5 mM the maximal [Cl^−^]_i_ change was augmented by addition of 107 AMPA synapses with a g_AMPA_ of 0.305 nS, while the [Cl^−^]_i_ decline at an initial [Cl^−^]_i_ of 25 mM was attenuated by this AMPA co-stimulation. (g) Statistical analysis of the voltage changes induced by simulated GDPs with the given initial [Cl^−^]_i_ and g_AMPA_. (h) Statistical analysis of [Cl^−^]_i_ changes induced by simulated GDPs with the given initial [Cl^−^]_i_ and g_AMPA_. Bars represent mean ± SD of 9 repetitions. Panel a used with permission from Lombardi et al. [17].

In a first set of simulations, we investigated the impact of g_AMPA_ on GABAergic [Cl^−^]_i_ changes during a simulated GDP. In accordance with previous observations [17,30], the massive GABAergic activity during a GDP led to substantial [Cl^−^]_i_ changes, with a [Cl^−^]_i_ increase by 3.94 ± 0.05 mM (n=9 repetitions) at a low initial [Cl^−^]_i_ of 5 mM and a [Cl^−^]_i_ decrease by -3.78 ± 0.08 mM (n=9) at an initial [Cl^−^]_i_ of 25 mM (black traces in Fig. 7e, f). Adding 107 AMPA synapses with a g_AMPA_ of 0.305 nS (i.e. the experimentally determined number and conductance of GABAergic inputs) led to a substantial increase in the activity-dependent [Cl^−^]_i_ transients to 4.95 ± 0.07 mM for an initial [Cl^−^]_i_ of 5 mM and significantly attenuated the activity-dependent [Cl^−^]_i_ decrease at 25 mM to -2.88 ± 0.06 mM (Fig. 7f red trace). A similar trend was also observed for other initial [Cl^−^]_i_ concentrations. Co-stimulation with 107 AMPA inputs increased the [Cl^−^]_i_ transients at low initial [Cl^−^]_i_ and attenuated the transients at high initial [Cl^−^]_i_ (Fig. 7h). Using a larger value for g_AMPA_ enhanced this impact on activity-dependent [Cl^−^]_i_ shift (Fig. 7f). At g_AMPA_ values of 30.5 and 305 nS at all investigated initial [Cl^−^]_i_, an increase in [Cl^−^]_i_ was induced (Fig. 7h), in accordance with the depolarized E_m_ induced by these conditions (Fig. 7g). In summary, these results indicate that physiological levels of AMPA co-stimulation can significantly affect the GABA_A_ receptor induced [Cl^−^]_i_ transients.

In the next series of simulations, we systematically altered the decay time constant τ_AMPA_ for all 107 AMPA receptor-mediated synaptic inputs during a GDP from the experimentally determined value of 11 ms to 5, 37 and 50 ms, while using g_AMPA_ of 0.305 nS and the experimentally determined values for the GABAergic synaptic inputs. Although this slight shift in τ_AMPA_ had only a minor effect on the size of the voltage deflections (Fig. 8a, c), it significantly modified the effect of AMPA co-stimulation on GABA_A_ receptor mediated [Cl^−^]_i_ (Fig. 8b, d). At an initial [Cl^−^]_i_ of 5 mM the enhancement of GDP-induced [Cl^−^]_i_ transient by AMPA (Δ_G_[Cl^−^]_i_) was reduced from 1.02 ± 0.06 mM (n=9 repetitions) at τ_AMPA_ = 11 ms to 0.48 ± 0.06 mM at τ_AMPA_ = 5 ms (Fig. 8b). Similarly, at an initial [Cl^−^]_i_ of 25 mM, Δ_G_[Cl^−^]_i_ was diminished by shortening τ_AMPA_ from 11 ms (0.93 ± 0.1 mM) to 5 ms (0.43 ± 0.07 mM) (Fig. 8b). In contrast, prolonging τ_AMPA_ to 37 or even 50 ms had a substantial impact on the time course and amount of the voltage deflections (Fig. 8a). Accordingly, the activity-dependent [Cl^−^]_i_ transients were systematically shifted toward more influx or less efflux, respectively (Fig. 8b-d). In summary, these results indicate that even slight changes in τ_AMPA_ can substantially influence the impact of AMPA co-activation on GABA_A_ receptor-mediated [Cl^−^]_i_ transients.

**Figure 8.**
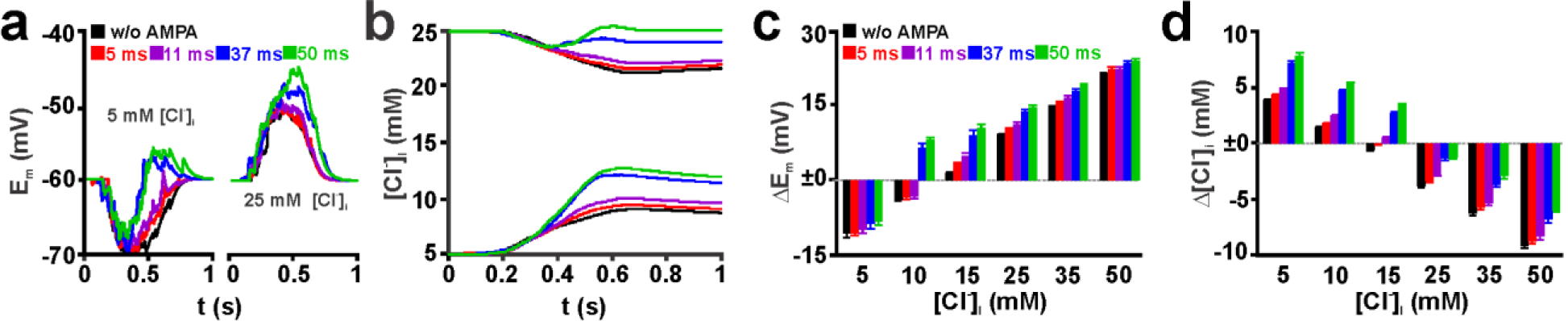
Effect of the decay time of AMPA receptor-mediated currents (τ_AMPA_) on activity-dependent [Cl^−^]_i_ transients induced by multiple spatially distributed synaptic inputs replicating GDP-like depolarizations. (a) Typical voltage traces in a simulated CA3 pyramidal neuron upon GABA stimulation (black trace) and GABA-AMPA co-stimulation with different τ_AMPA_. Left traces represent stimulations at initial [Cl^−^]_i_ of 5 mM, right traces at a [Cl^−^]_i_ of 25 mM. Note that a prolongation of τ_AMPA_ above 11 ms led to an obvious shift of the voltage response to positive values. (b) Average [Cl^−^]_i_ observed in these experiments. Note that the [Cl^−^]_i_ changes upon GABA-AMPA co-stimulation were only slightly shifted at τ_AMPA_ of 5 and 11 ms, while at 37 ms and 50 ms an obvious shift towards Cl^−^ efflux was induced. (c) Analysis of the voltage responses upon AMPA-GABA co-stimulation. (d) Analysis of the average [Cl^−^]_i_ changes upon GABA-AMPA co-stimulation. Note that for small initial [Cl^−^]_i_ longer τ_AMPA_ augmented the Cl^−^ -influx, while at high initial [Cl^−^]_i_ prolongation of τ_AMPA_ diminished the Cl^−^ -efflux. Bars represent mean ± SD of 9 repetitions.

### 2.5 Spatial and temporal constrains of AMPA-mediated infuences on GABAergic [Cl^−^]_i_ transients in a realistic neuronal model

Next we analyzed how the spatial relation between GABA and AMPA synapse activation influences the size of GDP-dependent [Cl^−^]_i_ transients. For this purpose, we first compared whether direct co-localization of each of the 107 AMPA synapses with a GABA synapse (100% spatial correlation) affects the AMPA-mediated shift in the GDP-dependent [Cl^−^]_i_ transients. These simulations revealed that the activity-dependent [Cl^−^]_i_ shifts with this 100% spatially correlated AMPA synapses were not significantly different from models with a random spatial distribution of AMPA synapses (Fig. 9a). Also a restriction of AMPA synapses in either the distal 25% or the proximal 25% of the dendrite length had no significant effect on the magnitude of activity-dependent [Cl^−^]_i_ transients (data not shown). We propose that this lack of an effect was caused by the fact that the overall density of GABA and AMPA synapses in each of the 56 dendrites was so high, that independent of the individual synaptic localization comparable effects on DF_GABA_ were induced. Therefore, we next located the GABA and AMPA synapses in different parts of the dendritic compartment. In these simulations the GABA_A_ receptor-induced [Cl^−^]_i_ shift without AMPA co-stimulation differ from the previous stimulations, as the Cl^−^-influx was restricted to a subset of dendrites (Fig. 9b). Under this strict regime, the location of the AMPA synapses had a slight effect on Δ_G_[Cl^−^]_i_. With all GABA synapses in the proximal dendrites and the AMPA synapses located in the distal dendrites a co-stimulation induced a Δ_G_[Cl^−^]_i_ of 0.33 ± 0.05 mM (n=9 repetitions) at an initial [Cl^−^]_i_ of 5 mM (Fig. 9b). In contrast, when all GABA synapses were located in the distal dendrites and all AMPA synapses in the proximal dendrites co-stimulation induced a Δ_G_[Cl^−^]_i_ of 0.52 ± 0.02 mM (Fig. 9b). A similar tendency was also observed for other initial [Cl^−^]_i_ (Fig. 9b). In summary, these results indicate that for a massive stimulation, like the modeled GDP, changes in the spatial correlation between AMPA and GABA receptors have only subtle effects on activity-dependent [Cl^−^]_i_ transients.

**Figure 9.**
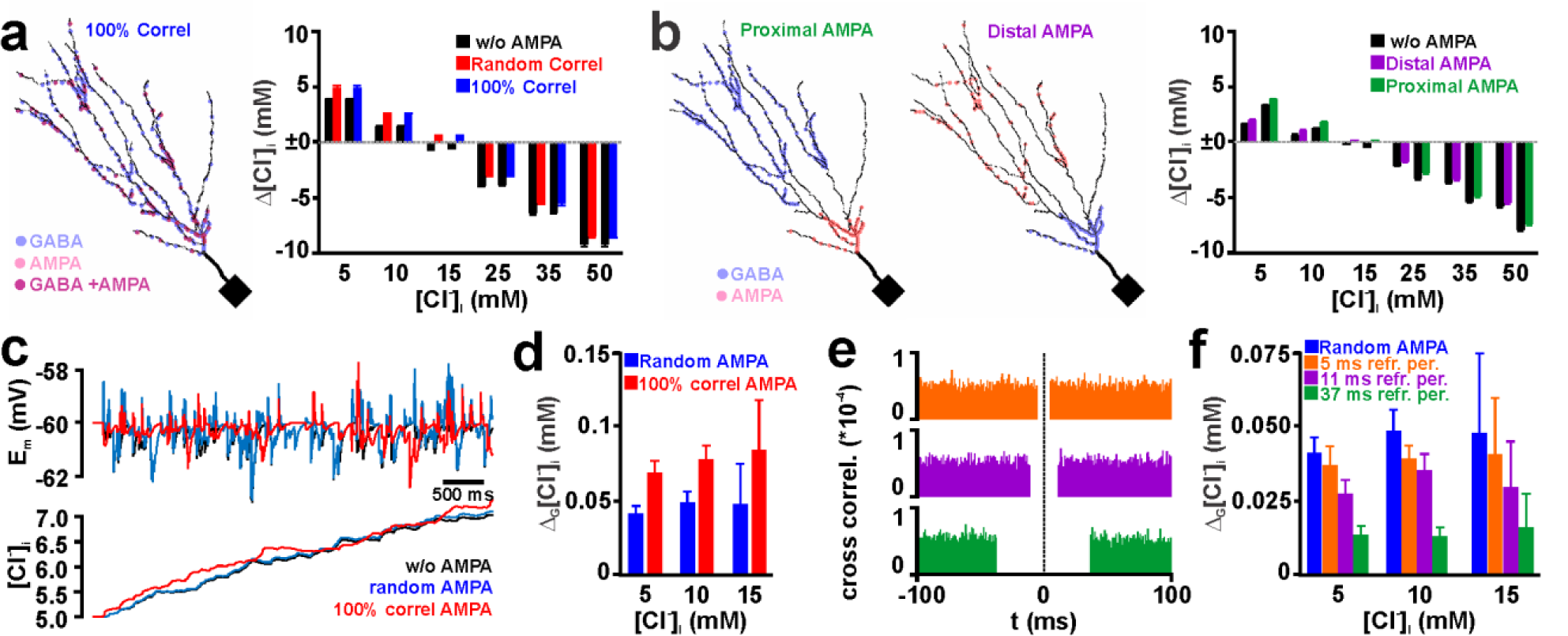
Effect of spatial and temporal correlation between multiple dendritic GABA and AMPA synapse activation on the activity-dependent [Cl^−^]_i_ transients. (a) Effect of randomly distributed (red columns) and GABA colocalized (blue columns) AMPA inputs on GDP-induced [Cl^−^]_i_ transients. Note that both distributions had identical effects. In the left panel the positions of the synapses in the dendritic compartment are plotted. (b) In the left panels the localization of AMPA receptors in proximal and distal dendrites are displayed. Note that GABA synapses were placed in the opposite (distal and proximal) compartment. The right plot indicates that significantly larger effects of AMPA co-stimulation were observed if the AMPA synapses were positioned in the proximal compartment. (c) Typical voltage and [Cl^−^]_i_ traces observed upon random stimulation of GABA synapses without AMPA co-stimulation (black), random AMPA co-stimulation (blue), or with temporally perfectly correlated AMPA inputs (red). (d) Statistical analysis of the additional [Cl^−^]_i_ increase upon AMPA co-stimulation (Δ_G_[Cl^−^]_i_). (e) Cross correlograms of simulations in which the AMPA stimuli were forced to a refractory period of 5 ms (orange), 11 ms (purple), and 37 ms (green). (f) Statistical analysis of Δ_G_[Cl^−^]_i_ at different refractory periods. Note that at refractory periods of ≥ 11 ms significantly smaller Δ_G_[Cl^−^]_i_ occurred. Bars represent mean ± SD of 9 repetitions.

Finally, we investigated the effect of the temporal correlation between AMPA and GABA synaptic inputs on the activity-dependent [Cl^−^]_i_ changes. Since it was not possible to generate a non-trivial decorrelation of AMPA and GABA synaptic inputs during GDP-like activity, we randomly stimulated 100 AMPA (g_AMPA_ = 0.305 pS/τ_AMPA_ = 11 ms) and 100 GABA (g_AMPA_ = 0.789 pS/τ_AMPA_ = 37 ms) synaptic inputs at a frequency of 20 Hz. While at an initial [Cl^−^]_i_ of 5 mM the random stimulation of only GABA synapses induced a [Cl^−^]_i_ increase by 2.06 ± 0.16 mM (n=9 repetitions), this increase was augmented by 0.04 ± 0.005 mM upon random co-stimulation of AMPA synapses (Fig. 9 c, d). Comparable Δ_G_[Cl^−^]_i_ values were also observed for initial [Cl^−^]_i_ of 10 mM and 15 mM. For higher initial [Cl^−^]_i_ the trend towards larger [Cl^−^]_i_ at AMPA co-stimulation maintained. However, the large variance in the individual maximal [Cl^−^]_i_ responses in each simulation and the resulting high SD values for Δ_G_[Cl^−^]_i_ values caused that these changes were not significant. If in this stimulation paradigm the AMPA stimuli occurred exactly at the same time points as the GABAergic inputs (i.e. 100% temporal correlation), Δ_G_[Cl^−^]_i_ was significantly increased to 0.065 ± 0.008 mM (at an initial [Cl^−^]_i_ of 5 mM), to 0.078 ± 0.009 mM (at 10 mM [Cl^−^]_i_), and to 0.084 ± 0.034 mM (at 15 mM [Cl^−^]_i_) (Fig. 9d).

To investigate the effect of a decorrelation between GABA and AMPA inputs, we implemented an algorithm that generates refractory periods of 5, 11, and 37 ms around each GABAergic stimulus in which AMPA inputs are omitted (Fig. 9e). These experiments revealed that for an initial [Cl^−^]_i_ of 5 mM the Δ_G_[Cl^−^]_i_ values are significantly decreased from 0.041 ± 0.005 mM (n=9 repetitions at random correlation) to 0.027 ± 0.004 at a refractory period of 11 ms, and to 0.014 ± 0.003 at a refractory period of 37 ms (Fig. 9f). At a refractory period of 5 ms Δ_G_[Cl^−^]_i_ decreased only slightly and not significantly to 0.037 ± 0.006. A comparable reduction was also observed for an initial [Cl^−^]_i_ of 10 mM (Fig. 9f). At 15 mM only for a refractory period of 37 ms a significant reduction in Δ_G_[Cl^−^]_i_ was observed (from 0.048 ± 0.026 mM to 0.016 ± 0.011 mM). In summary, these results support the finding that the temporal correlation between GABA and AMPA synapses enhance activity-dependent [Cl^−^]_i_ shifts, in particular if AMPA and GABA inputs are correlated within the time range of their decay time constants.

## 3. Discussion

Recent studies provide increasing evidence that GABAergic responses show an activity- and compartment-dependent behavior due to ionic plasticity in [Cl^−^]_i_ [11,24]. Here we used realistic biophysical modeling to study how coincident glutamatergic inputs enhance the [Cl^−^]_i_ changes induced by GABAergic activation. The main findings of this computational study can be summarized as follows: 1.) Glutamatergic co-stimulation had a direct effect on the GABAergic [Cl^−^]_i_ changes, thereby enhancing Cl^−^ influx at low initial [Cl^−^]_i_ and attenuating or even reversing the Cl^−^ efflux caused at high initial [Cl^−^]_i_. 2.) Massive glutamatergic co-stimulation promotes Cl^−^ influx at all initial [Cl^−^]_i_, whereas physiological levels of glutamatergic co-stimulation mediate biphasic [Cl^−^]_i_ changes at intermediate [Cl^−^]_i_ levels typical for immature neurons. 3.) Keeping the decay kinetics of glutamatergic inputs below that of GABAergic inputs attenuated the [Cl^−^]_i_ changes. 4.) The spatial and temporal correlation between glutamatergic and GABAergic inputs has a stringent influence on the [Cl^−^]_i_ changes, with a surprisingly large temporal interval with correlation-independent [Cl^−^]_i_ changes. 5.) In a realistic model of physiologically relevant synaptic activity that replicates GDPs the conductance and kinetics of correlated glutamatergic activity has a substantial impact on the [Cl^−^]_i_ changes. 6.) While the spatial correlation between distributed GABA and glutamatergic synapses has only a minor effect on activity-dependent [Cl^−^]_i_ changes, their temporal correlation has strong effects.

In this study we investigated the effect of co-activation of glutamatergic AMPA receptors on GABA receptor-dependent [Cl^−^]_i_ changes over a wide range of initial [Cl^−^]_i_, which gave us the opportunity to evaluate the role of ionic plasticity for mature and developing nervous systems [3,40]. In the mature brain GABAergic activity causes an increase in [Cl^−^]_i_ [11,16,25,41,42], thereby decreasing the inhibitory action of the GABAergic system [13,26,27,33]. The present study demonstrates that glutamatergic co-stimulation enhances the GABAergic Cl^−^-influx and thus enhances the activity-dependent [Cl^−^]_i_ increase, as has recently been shown in-vitro [32] and in-silico [33]. As a consequence, the inhibitory action of GABA may be even more impaired upon a co-activation of glutamate receptors [33]. The high basal [Cl^−^]_i_ in immature neurons leads to GABAergic Cl^−^-efflux and thus to a decrease in [Cl^−^]_i_ upon GABAergic activation [18,43]. This [Cl^−^]_i_ decline can be associated with a loss of excitatory action and/or an enhancement of the shunting inhibitory effect of GABA receptors [21,44,45]. The present study demonstrates for the first time that in immature neurons a co-activation of GABA and glutamate receptors attenuates the activity-dependent [Cl^−^]_i_ decrease, thereby stabilizing the depolarizing actions of GABA.

The absolute values of the additional [Cl^−^]_i_ changes induced by AMPA co-stimulation in our ball-and-stick simulations are small, however it has to be taken into account that these changes represent the [Cl^−^]_i_ changes induced at a single synapse with a single synaptic stimulus. Physiological levels of synaptic activity may thus cause significantly larger [Cl^−^]_i_ changes, as has been shown in-vitro [32]. Even physiological levels of glutamatergic co-stimulation (0.305 - 3.05 nS, representing 1-10x spontaneous synaptic inputs) mediate in our model a reliable increase in Cl^−^ influx at low initial [Cl^−^]_i_, implicating that glutamatergic co-stimulation will enhance ionic plasticity in mature neurons. Thereby coincident glutamatergic activity provides an additional challenge on the [Cl^−^]_i_ homeostasis in mature neurons and can substantially contribute to an activity-dependent decline in the inhibitory action of GABAergic synapses [33,46,47]. But even small Cl^−^ changes can cause substantial functional alterations for neuronal computation, as has been recently demonstrated in-silico [48,49]. Subtle changes in GABAergic membrane responses may also affect the temporal coding of synaptic inputs [44] or directly interfere with the action potential threshold [50].

As mentioned before, the high [Cl^−^]_i_ in immature neurons and the resulting GABAergic Cl^−^-efflux results in a condition in which glutamatergic co-stimulation attenuates GABAergic [Cl^−^]_i_ loss and thereby stabilizes depolarizing GABAergic actions. In addition, our computational model demonstrates that at [Cl^−^]_i_ values between 15 and 35 mM, which are in the typical range determined in immature neurons [4,51,52], moderate glutamatergic co-stimulation (0.305 - 3.05 nS) caused biphasic [Cl^−^]_i_ changes. These biphasic [Cl^−^]_i_ changes rely on the fact that the activation of glutamatergic receptors transiently pushes the membrane potential above E_Cl_ and thus supports a Cl^−^-influx during this interval. The biphasic [Cl^−^]_i_ changes reduce the maximal [Cl^−^]_i_ decline at the end of the GABAergic postsynaptic potential. By this process the activity-dependent [Cl^−^]_i_ decline as well as the burden of glutamatergic co-stimulation on [Cl^−^]_i_ homeostasis is reduced. It is tempting to speculate that this limitation of activity-dependent Cl^−^-fluxes may be one reason for the inefficient Cl^−^-transport in the immature CNS [4,22]. The functional implications of activity dependent [Cl^−^]_i_ changes in immature neurons are harder to predict, as depolarizing GABAergic responses can mediate inhibitory [53,54] as well as excitatory actions [55–57]. However, it can be estimated that an attenuation of the GABAergic [Cl^−^]_i_ depletion will prevent/ameliorate a loss of GABAergic excitation and will prevent/ameliorate an increase in the inhibitory effect of depolarizing GABAergic responses. Thereby coincident glutamatergic activity, which is a typical feature of the correlated network events in developing neuronal systems [36,58], can stabilize GABAergic functions.

Our simulation demonstrated that τ_AMPA_ has, as expected, a substantial influence on Δ_G_[Cl^−^]_i_, with faster AMPA events reducing the amount of ionic plasticity. Thus the shorter AMPA receptor-mediated responses at adult synapses [59,60] can reduce the impact of AMPA/GABA co-activation on [Cl^−^]_i_ homeostasis, in addition to making transmission more precise. In this respect, it is also tempting to speculate that conditions which permit the unblocking of NMDA-receptors (a subtype of glutamate receptors that is characterized by a rather long decay and that plays an essential role in learning, [61]), will also lead to more massive burden on [Cl^−^]_i_ homeostasis. The resulting short-term decline in GABAergic inhibition will make neurons more prone to the induction of long-term potentiation.

In addition to this expected influence of τ_AMPA_ on ionic plasticity, our simulations revealed a complex dependency of Δ_G_[Cl^−^]_i_ on τ_GABA_ and τ_AMPA_. Interestingly the maximal Δ_G_[Cl^−^]_i_ values were obtained when τ_GABA_ was slightly larger than τ_AMPA_. An additional prolongation of τ_GABA_ even reduced Δ_G_[Cl^−^]_i_ and thus diminished ionic plasticity. Intriguingly, such a prolongation of τ_GABA_ led to the appearance of a “plateau”-like phase, in which Δ_G_[Cl^−^]_i_ is insensitive to the latency between GABA and AMPA inputs. The analysis of the Cl^−^-fluxes in the phase diagram revealed that this “plateau”-like phase was due to the fact, that the Cl^−^-fluxes converged over a wide range of AMPA/GABA latencies in the trajectory of [Cl^−^]_i_ changes obtained by synchronous GABAergic and glutamatergic stimulation. For short τ_AMPA_ no “plateau”-like phase could be observed because under these conditions the time course of E_m_ was mainly determined by the membrane time constant. This explanation was supported by our observation that an artificial increase in the membrane time constant by an augmented C_m_ caused a dissipation in the phase plane plot from two attractors towards more dissociated trajectories. For larger τ_AMPA_ of 37 and 50 ms such a “plateau”-line phase was not reached because at this longer time constant the decay of the voltage deflection is determined by both the inactivation of AMPA- and GABA_A_ receptors and thus a gradual decrease in DF_GABA_ occurred during the course of co-activation.

It has been suggested that under mature conditions ionic plasticity of [Cl^−^]_i_ can act as a coincidence detector for simultaneous GABAergic and glutamatergic synaptic inputs [32]. Via such a coincidence detection GABAergic inhibition at will be particularly attenuated by coincident glutamatergic inputs. The resulting [Cl^−^]_i_ increase and the corresponding reduction in the inhibitory GABAergic capacity will subsequently promote the relay of excitation in neuronal networks. The intriguing finding of the present manuscript, that the effect of glutamatergic co-stimulation on the GABAergic [Cl^−^]_i_ increase was stable for a considerable latency interval between GABAergic and glutamatergic inputs would allow the system to use a stable mechanism for adjusting the gating of excitatory information. In this respect, the striking asymmetry of in Δ_G_[Cl^−^]_i_ under physiological conditions of the decay times (τ_AMPA_ < τ_GABA_) implies a spike-time dependency of ionic plasticity. While glutamatergic inputs preceding GABAergic inputs have only a small effect on Δ_G_[Cl^−^]_i_, a stable shift in Δ_G_[Cl^−^]_i_ was induced when glutamatergic synapses were activated during the decay phase of GABAergic responses. A possible functional implication of this spike-time dependency would be that common feed-forward inhibitory circuits, in which synaptic inhibition always occurs after the glutamatergic inputs, will be only minimally affected by ionic plasticity, guaranteeing an efficient and stable feed-forward inhibition. Only in cases in which glutamatergic excitation was induced during ongoing GABAergic stimulation, a substantial ionic plasticity would operate. By this mechanism the inhibitory capacity of GABA will be reduced particularly in the postsynaptic neurons that receive frequent glutamatergic inputs, thereby facilitating particularly these pathways.

In summary, our results demonstrate that glutamatergic co-activation has a prominent time- and space-dependent effect on the amount of GABA-receptor mediated [Cl^−^]_i_ changes. This ionic plasticity depends on the properties of glutamatergic inputs and their temporal correlation with the GABAergic inputs. These glutamatergic modulations of GABAergic ionic plasticity can contribute to short-term memory and may profoundly influence information processing in the developing and mature nervous system.

## 4. Materials and Methods

### Compartmental modeling

The biophysically realistic compartmental modeling was performed using the NEURON environment (neuron.yale.edu). The source code of models and stimulation files used in the present paper can be found in ModelDB (http://modeldb.yale.edu/266765; reviewer password is []). For compartmental modelling we used in the first simulations a simple ball and stick model (soma with d=20 µm, linear dendrite with l=200 µm, diameter 1 µm, and 103 nodes; *cell_soma_dendrite*.*hoc*). If not otherwise noted the dendrite was electrically and diffusionally detached from the soma to analyse dendritic [Cl^−^]_i_ transients (*cell_isolated_dendrite*.*hoc*). In further experiments we used a reconstructed CA3 pyramidal cell (*Cell1_Cl-HCO3_Pas*.*hoc*; [17]). This reconstructed neuron contained a soma (d= 15 µm), a dendritic trunk (d= 2 µm, l = 32 µm, 9 segments) and 56 dendrites (d= 0.36 µm, 9 segments each). In all of these compartments a specific axial resistance (R_a_) of 34.5 Ωcm and a specific membrane capacitance (C_m_) of 1 µFcm^-2^ were implemented. C_m_ was varied in two experiments to accelerate/decelerate the membrane time constant. A specific membrane conductance (g_*pas*_) of 0.5 mS/cm^2^ with a reversal potential of -60 mV was inserted in all neuronal elements.

GABA_A_ synapses were simulated as a postsynaptic parallel Cl^−^ and HCO_3^-^_ conductance with exponential rise and exponential decay:

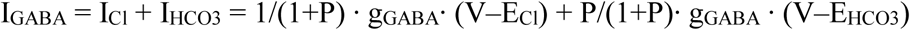

where P is a fractional ionic conductance that was used to split the GABA_A_ conductance (g_GABA_) into Cl^−^ and HCO_3^-^_ conductance. E_Cl_ and E_HCO3_ were calculated from the Nernst equation. The GABA_A_ conductance was modeled using a two-term exponential function, using separate values of rise time (0.5 ms) and decay time (variable, mostly 37 ms). Parameters used in our simulations were as follows: [Cl^−^]_o_ = 133.5 mM, [HCO3^-^]_i_ = 14.1 mM, [HCO3^-^]_o_ = 24 mM, temperature = 31^°^C, P = 0.18. The conductance values for GABA synapses varied as given in the main text. AMPA synapses were modeled by an Exp2Syn point process using a reversal potential of 0 mV, a tau 1 value of 0.1 ms and a tau2 value of 11 ms, in accordance with the experimentally determined value [17], except where noted. The conductance values for AMPA synapses varied as given in the main text.

For the ball and stick model a single GABA_A_ synapse was placed in the middle of the dendrite, except where noted. The AMPA synapse was placed as indicated in the figure legends. For the simulation of a GDP in the reconstructed CA3 neuron 534 GABAergic synapses and 107 AMPA synapses were randomly distributed within the dendrites of the reconstructed neuron, except where noted. The number of GABA and AMPA synapses replicate the estimated synapse numbers during experimentally recorded GDPs [17]. GABA and AMPA inputs were activated stochastically using a normal distribution (µ = 600ms, σ = 9000 ms for GABA and µ = 650ms, σ = 8500 ms for AMPA) that emulates the distribution of GABAergic and glutamatergic PSCs during a GDP observed in immature hippocampal CA3 pyramidal neurons [17]. The properties of these synapses were given in the results part and/or the corresponding figure legends. Repetitions of these simulations were generated by altering the seed values for the random functions between 1 and 9.

For the modeling of the GABA_A_ receptor-induced [Cl^−^]_i_ and [HCO_3_ ^-^]i changes, we calculated ion diffusion and uptake by standard compartmental diffusion modeling. To simulate intracellular Cl^−^ and HCO3^-^ dynamics, we adapted our previously published model [25,30]. Longitudinal Cl^−^ diffusion along dendrites was modeled as the exchange of anions between adjacent compartments. For radial diffusion, the volume was discretized into a series of four concentric shells around a cylindrical core and Cl^−^ or HCO_3_^-^ was allowed to flow between adjacent shells. The free diffusion coefficient of Cl^−^ inside neurons was set to 2 µm^2^/ms [14]. Since the cytoplasmatic diffusion constant for HCO_3_^-^ is, to our knowledge, unknown, we also used a value of 2 µm^2^/ms. To simulate transmembrane transport of Cl^−^ and HCO_3_^-^, we implemented an exponential relaxation process for [Cl^−^]_i_ and [HCO_3_^-^]_i_ to resting levels [Cl^−^]_i_^rest^ or [HCO ^-^] ^rest^ with a time constant τ_Ion_.

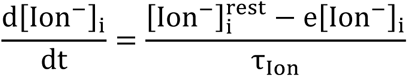

Cl^−^ transport was in most experiments (if not otherwise noted) modeled as bimodal process, for [Cl^−^]_i_ < [Cl^−^]_i_^rest^ τ_Ion_ was set to 174 s to emulate an NKCC1-like Cl^−^ transport mechanism. For [Cl^−^]_i_ > [Cl^−^]_i_^rest^ τ_Ion_ was set to 321 s to emulate passive Cl^−^ efflux (both values obtained from unpublished experiments on immature rat CA3 hippocampal neurons).

The impact of GABAergic Cl^−^ currents on [Cl^−^]_i_ and [HCO ^-^] was calculated as:

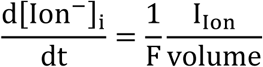

To simulate the GABAergic activity during a GDP, a unitary peak conductance of 0.789 nS and a decay of 37 ms were applied to each GABAergic synapse, in accordance with properties of spontaneous GABAergic postsynaptic currents in CA3 pyramidal neurons [17].

For the isolated neurons the [Cl^−^]_i_ concentration was averaged over all segments of the dendrite, except where noted. For the simulated neurons we analyzed mean [Cl^−^]_i_ of all dendrites:

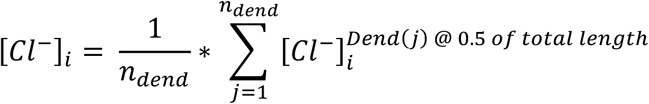

This procedure mimics the experimental procedure of Lombardi et al. [17], who determined E_GABA_ by focal application in the dendritic compartment.

For the calculation of Δ[Cl^−^]_i_, the maximal deviation of [Cl^−^]_i_ upon a GABAergic stimulus ([Cl^−^]_i_^S^) was subtracted from the resting [Cl^−^]_i_ before the stimulus ([Cl^−^]_i_^R^). For biphasic responses both minimal and maximal [Cl^−^] ^R^ were determined and displayed. In some cases, the manifest Δ[Cl^−^]_i_ for these responses was calculated as:

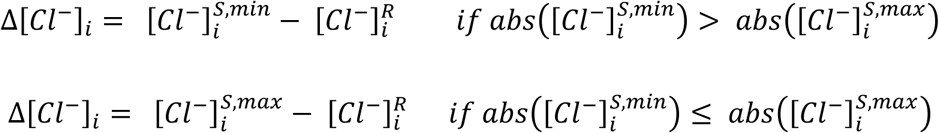

To quantify the influence of AMPA co-stimulation on the GABAergic [Cl^−^]_i_ transients, the amount of additional [Cl^−^]_i_ changes induced by AMPA co-stimulation (Δ_G_[Cl^−^]_i_) was calculated from the Δ[Cl^−^]_i_ values with/without AMPA co-stimulation as follows:

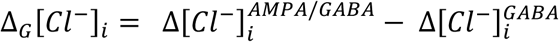

## Acknowledgements

The authors thank Beate Krumm for her excellent technical support. This research was funded by grants of the Deutsche Forschungsgemeinschaft to WK (KI-835/3) and to HJL (CRC 1080, A01), by the BMBF grant to PJ (No. 031L0229) and by funds from the von Behring Röntgen Foundation to PJ.

## Abbreviations

DF_GABA_: Electromotive driving force on GABAergic currents
E_Cl_: Equilibrium potential for Cl^−^
E_GABA_: Reversal potential of GABAergic currents
E_HCO3_: Equilibrium potential for HCO_3_^-^
E_m_: Membrane voltage
GABA: γ-Amino butyric acid
GDP: Giant depolarizing potential
g_GABA_: Conductance of GABAergic synapse
g_pas_: Passive membrane conductance
KCC: K^+^-Cl^−^-Cotransporter
NKCC1: Na^+^-K^+^-Cl^−^-Cotransporter, Isoform 1
n_GABA_: Number of GABAergic synapses
P_HCO3_: Relative HCO_3_^-^ permeability of GABA_A_ receptors
V_Hold_: Holding potential
τ_Cl_: Time constant of [Cl^−^] relaxation
τ_GABA_: Decay time constant of GABA_A_ receptors
τ_HCO3_: Time constant of [HCO_3_^-^] relaxation

